# The hormone Fibroblast Growth Factor 19 stimulates water intake

**DOI:** 10.1101/2021.10.26.466031

**Authors:** José Ursic-Bedoya, Carine Chavey, Lucy Meunier, Guillaume Desandré, Anne-Marie Dupuy, Iria Gonzalez-Dopeso Reyes, Thierry Tordjmann, Eric Assénat, Urszula Hibner, Damien Gregoire

## Abstract

Fibroblast growth factor 19 (FGF19) is a hormone with pleiotropic metabolic functions, leading to ongoing development of analogues for the treatment of metabolic disorders. On the other hand, FGF19 is overexpressed in a sub-group of hepatocellular carcinoma (HCC) patients and has oncogenic properties. It is therefore crucial to precisely define FGF19 effects, notably chronic exposure to elevated concentrations of the hormone. We used hydrodynamic gene transfer approach to generate a transgenic mouse model with long-term FGF19 hepatic overexpression. Here we describe a novel effect of FGF19, namely stimulation of water intake. This phenotype, lasting at least over a 6-month period, depends on signaling in the central nervous system and is independent of FGF21, although it mimics some of its features. We further show that HCC patients with high levels of circulating FGF19 have a reduced natremia, indicating dispogenic features. The present study provides evidence of a new activity for FGF19, which could be clinically relevant in the context of FGF19 overexpressing cancers and treatment of metabolic disorders by FGF19 analogues.

## Introduction

Fibroblast growth factor 19, FGF19, (and its mouse ortholog, FGF15), functions as a hormone, notably controlling bile acids synthesis and nutrient metabolism (Angelin *et al*, 2012). FGF15/19 has been reported to exert multiple effects in the regulation of glucose and lipid metabolism, as well as in energy expenditure and body adiposity, through its actions on liver, muscle, white adipose tissue and brain (Gadaleta & Moschetta, 2019).

Under physiological conditions, FGF19 is expressed during the postprandial phase by enterocytes of the terminal ileum(Inagaki *et al*, 2005). The expression by hepatocytes has also been described in cholestasis (Schaap *et al*, 2009; Wunsch *et al*, 2015; Hasegawa *et al*, 2019). FGF19 belongs to the endocrine FGF subfamily, along with FGF21 and FGF23. Endocrine-FGF lack the heparin binding domain found in canonical FGFs and act through the binding to a FGF receptor (FGFR 1 to 4) in complex with a co-receptor, β-klotho (KLB) for FGF19 and FGF21 (Suzuki *et al*, 2008; Wu *et al*, 2010). While many of FGF19 activities are thought to be mediated by FGFR4/KLB signaling, FGF19 and FGF21 also display overlapping metabolic regulations through activation of FGFR1c/KLB signaling (Wu *et al*, 2011), notably in the hypothalamus (Lan *et al*, 2017; Perry *et al*, 2015). Because of their metabolic effects, FGF19 and FGF21 pathways are considered as promising therapeutic targets for several diseases (Degirolamo *et al*, 2016). In particular, FGF19 has generated a great interest in pharmacological research for treatment of chronic liver diseases such as nonalcoholic steatohepatitis (Harrison *et al*, 2018, 2020), primary biliary cholangitis (Mayo *et al*, 2018) and primary sclerosing cholangitis (Hirschfield *et al*, 2019).

On top of its metabolic effects, FGF19/FGFR4 pathway shows oncogenic functions in several cancers, notably hepatocellular carcinoma, the most frequent primary liver cancer. The 11q13.3 genomic region containing FGF19 is frequently (4-15%) amplified in HCC tumors (Sawey *et al*, 2011; Wang *et al*, 2013; Schulze *et al*, 2015), and tumor cells overexpression of FGF19 independently of amplification has also been reported (Caruso *et al*, 2019; Hatlen *et al*, 2019). Moreover, HCC patients display higher circulating levels of FGF19 than control population (Maeda *et al*, 2019). These findings led to the development of potent and selective FGFR4 inhibitors, as well as monoclonal antibodies (Kim *et al*, 2019; Bartz *et al*, 2019), which are currently tested in clinical trials for FGF19-driven hepatocarcinoma, underscoring the therapeutic promise of targeting the FGF19 pathway in HCC (Gadaleta & Moschetta, 2021).

Because of these paradoxical effects on metabolism and oncogenesis, it is therefore crucial to better delineate the complex interplay of FGF19 roles and actions. In this study, we generated a mouse model of FGF19 overexpression by hepatocytes, and identified a new metabolic effect of FGF19, namely stimulation of water intake. We further report that HCC patients with increased FGF19 circulating concentrations show features of disturbance of the hydrosodic balance.

## Results and Discussion

### Mouse model of FGF19 expression

We used hydrodynamic gene transfer (HGT) technique (Liu *et al*, 1999; Zhang *et al*, 1999) to generate a mouse model in which FGF19 is overexpressed by a fraction of hepatocytes. The rapid injection of a large volume of solution (10% w/v) in the tail vein leads to *in vivo* transfection of hepatocytes. Combination of plasmids encoding the Sleeping Beauty transposase SB100X and FGF19 was used to establish stable expression of the human hormone (Figure 1A). We routinely detect by epifluorescence microscopy one to two percent of transfected hepatocytes, which persisted up to 6 months after the injection (Figure 1B). The long-term expression of the transgene was confirmed by RT-qPCR analysis (Figure 1C). The level of FGF19 in plasma, detected by ELISA, was around 10 ng/ml (CI_95%_= [3.5-15.0]) two weeks post-transfection and remained high, despite its gradual decrease, to reach 1 ng/ml at 6 months post-injection (CI_95%_= [0.76 -1.11]) (Figure 1D). These concentrations are supra-physiological, as healthy human subjects display circulating concentrations of 50-590 pg/mL (Angelin *et al*, 2012); however, they are close to those observed in a subset of HCC patients (Maeda *et al*, 2019). Of note, these pathologically relevant concentrations are 100 to 1000-fold inferior to those reported in a model of AAV infection (1 μg/mL (Zhou *et al*, 2017)). FGF19-expressing mice showed the previously characterized transcriptional repression of *Cyp7a1* and *Cyp8b1*, key enzymes of bile acids synthesis from cholesterol (Figure 1E). Serum 7a-hydroxy-4-cholesten-3-one (C4) level, a biomarker of bile acid synthesis, was also significantly reduced on FGF19-expressing mice (Figure 1E, 10.6 ng/mL CI_95%_= [4.9; 16.2] for controls *vs* 1.3 ng/mL CI_95%_= [0.8; 1.8] for FGF19, p=0.0027). As previously reported, FGF19 exerted a significant effect on the body weight (Figure 1F) (Lan *et al*, 2017), while histological analysis of livers revealed no visible effect of FGF19 overexpression (Figure 1G). Altogether, these results indicate that the secreted FGF19 is active, and that the circulating levels are compatible with its physiological functions.

**Figure 1:**
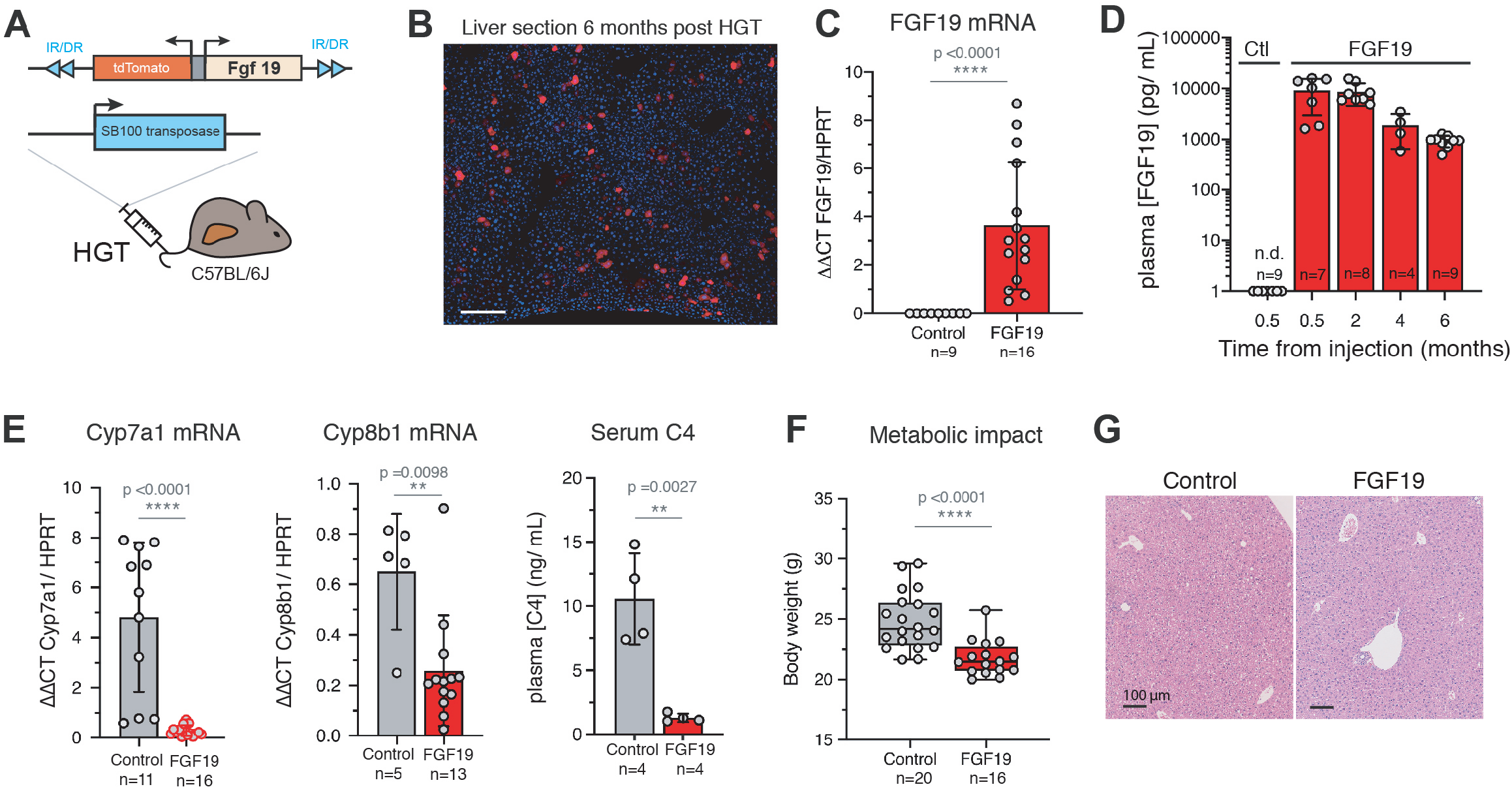
*in vivo* stable transfection of hepatocytes with FGF19 leads to long term secretion of biologically active FGF19. **A**. Mouse model strategy based on hydrodynamic gene transfer (HGT). **B**. Cryosection of liver 6 months after HGT, showing stably transfected hepatocytes that express TdTomato. Scale bar: 100μm. **C**. qPCR quantification of FGF19 mRNA in control and FGF19 transfected livers 6 months post injection. **D**. Plasma levels of FGF19 detected by ELISA. Plasmids combined with transposase are indicated, control-RFP («Ctl») and FGF19-RFP («FGF19») respectively. The number of mice analyzed for each time point is indicated. **E**. qPCR quantification of mRNA expression of Cyp7a1 and Cyp8b1, target genes of FGF19/FGFR4 pathway and serum 7 α-hydroxy-4-cholesten-3-one (C4) levels, serum biomarker of bile acid synthesis. **F**. Body weight of control and FGF19 HGT mice at 6 months. **G**. HES staining of liver sections of control and FGF19 HGT mice after 6 months, showing no visible alteration on liver morphology. Data information: In (C-F): data are presented as mean ± SD. * ps:0.05, **p <0.01, *** p<0.001, **** p<0.0001. C-E: Mann-Whitney test, F: unpaired Student’s t-test. N.s: not significant. n.d: not detectable.

### FGF19 overexpression increases water intake

While studying metabolic effects in FGF19 transgenic mice, we serendipitously observed that their cages were very wet compared to those of control mice, suggesting increased urine production. This prompted us to measure the animals’ water intake. We observed that FGF19 expressing mice drank 6.6 to 7.8 mL/day (CI_95%_ of the mean) compared to 2.9 to 3.1 mL/day for control animals hydrodynamically injected with an empty vector (p-value <0.0001) (Figure 2A). Strikingly, this phenotype was stable in time for at least 5 months. The increase of water intake was associated with a significant decrease of urine osmolality (Figure 2B, 1744 mOsm/kg/H20 CI_95%_= [1512; 1976] for controls *vs* 723 mOsm/kg/H20 CI_95%_= [568; 879] for FGF19, p<0.0001). We then measured urine production in individual metabolic cages and found that FGF19^+^ mice have an increased urine output compared to controls (Figure 2C, 490 μL/day CI_95%_= [204; 775] for controls *vs* 2871 μL/day CI_95%_= [1589; 4153] for FGF19, p=0.0004) and confirmed the phenotype of decreased urine osmolality (Figure 2D). Altogether, these results indicate that FGF19 overexpression mimics a phenotype of diabetes insipidus.

**Figure 2:**
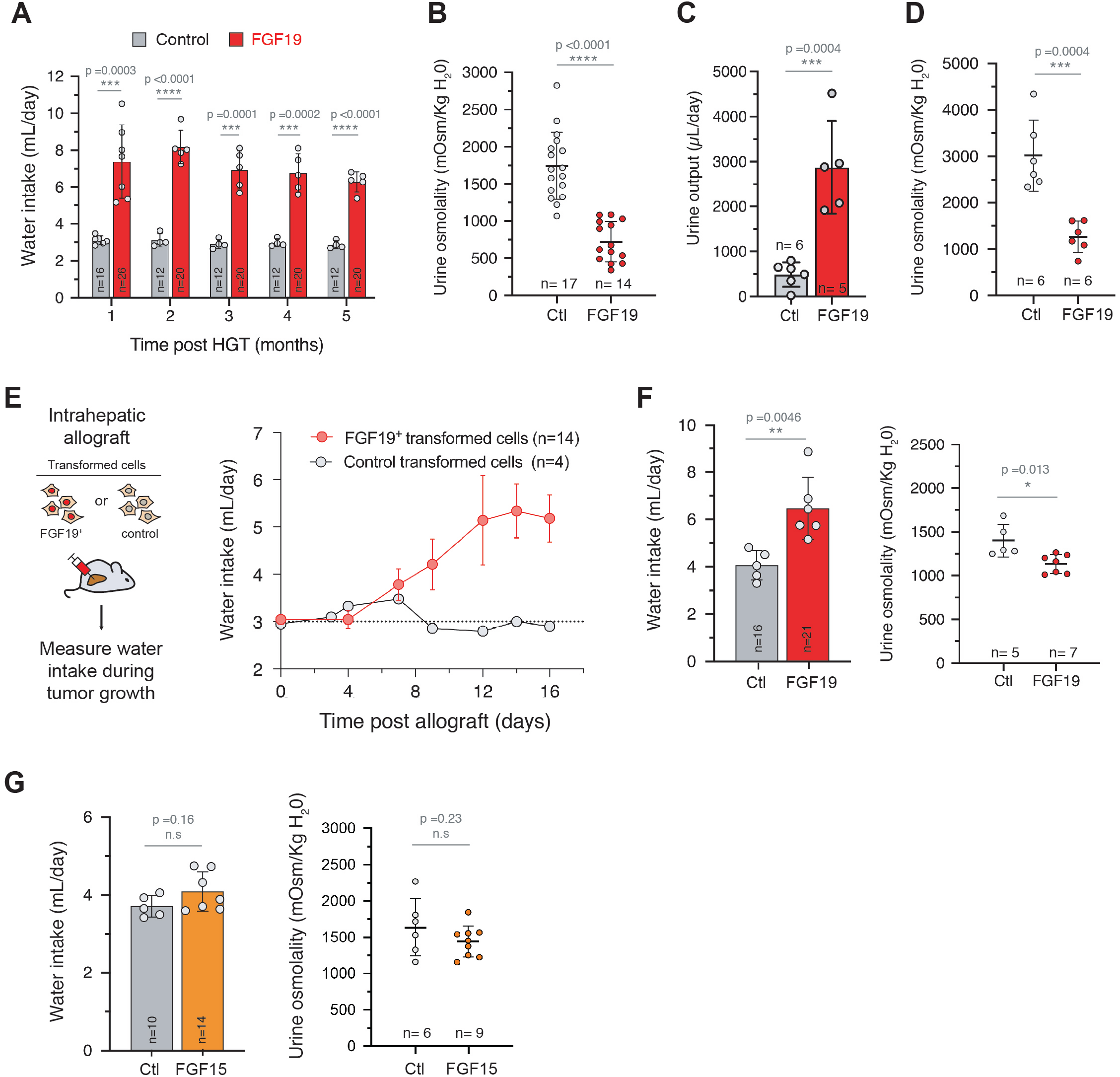
FGF19 increases water intake and urine output. **A**. Water consumption of control or FGF19+ female mice. The number of mice analyzed for each time point is indicated and each dot represents the mean value of an independent cage over one month. **B**. Urine osmolality of control and FGF19+ female mice at 3 months post HGT. **C**. Daily urine output in female mice housed in metabolic cages. **D**. Urine osmolality of control and FGF19+ female mice housed in metabolic cages. **E**. Water consumption of female mice following intrahepatic allograft of FGF19+ (MYC-sgTp53-FGF19) and control (MYC-sgTp53) cell lines. Mean water consumption is shown, corresponding to 14 mice (3 cages) or 4 mice (1 cage). Dotted line represents the mean consumption of C57BI/6J mice on the same period in the animal facility (n=15). **F**. Effect of FGF19 overexpression on water intake and urine osmolality on male mice 1 month post HGT. **G**. Water consumption and urine osmolality offemale mice injected with FGF15-BFP (FGF15) or BFP plasmid (control, Ctl) 1 month post HGT. Data information: In (A-G): data are presented as mean± SD. * p≤0.05, ** p<0.01, *** p<0.001, **** p<0.0001. n.s: not significant. For water intake mesures, each dot represents the mean value of an independent cage with two to five mice. For all panels, unpaired t-test statistical significance is indicated.

Next, we assessed the effect of FGF19 overexpression on water drinking in another experimental system, in which FGF19 is produced by hepatic tumor cells. To model FGF19-negative and FGF19-positive hepatic tumors, we realized intrahepatic allografts of oncogenic cell lines derived from Myc-sgTrp53 or Myc-sgTrp53-FGF19 tumors, both driven c-Myc oncogene in conjunction with inactivation of the p53 tumor suppressor, associated for the latter with FGF19 expression. Whereas Myc-sgTrp53 tumor development had no impact on water intake, mice injected with FGF19-expressing cells showed progressive rise of water intake from 3 to 7 mL/day as tumors grew (Figure 2E). Mean circulating FGF19 levels at 3 weeks in mice injected with Myc-sgTp53-FGF19 cell were 153 ng/mL (CI_95%_= [81; 225]), close to those obtained with AAV-driven FGF19 liver overexpression(Zhou *et al*, 2017). Thus, FGF19-driven increase of water intake by FGF19 was confirmed in a second experimental model. Importantly, this phenotype might be relevant for the subset of HCC patients overexpressing FGF19. It is unclear why the drinking phenotype has not been previously described in FGF19 overexpressing mice (Zhou *et al*, 2017; Nicholes *et al*, 2002). We note however that it would have probably escaped our notice if we were not directly handling the cages.

To determine if the diabetes insipidus phenotype was sex dependent, we next overexpressed FGF19 by HGT in C57Bl/6J males. Again, the FGF19 expression increased water intake, leading to a mean water consumption of 6.5 mL/ day (CI_95%_ = [5.1; 7.9]) during the first month after HGT (Figure 2F). We next investigated whether the overexpression of FGF15, the FGF19 mouse ortholog, also gave rise to the water drinking phenotype. Hydrodynamic gene transfer of FGF15 encoding plasmid did not significantly increase water intake nor reduced urine osmolality (Figure 2G). FGF15 shares only 53% of the amino acid sequence with FGF19 (Nishimura *et al*, 1999); notably, FGF15 has an unpaired cysteine residue (Cys-135) that forms an intermolecular disulfide bridge to structure a homodimer. It has been suggested that this particularity leads to a diminished receptor activation and metabolic effects compared to FGF19 (Zhou *et al*, 2017).

### FGF19 effect on the central nervous system

To test if the role of FGF19 in water homeostasis is central or peripheral, we performed an intraperitoneal injection to subject mice to an acute increase in water load. Cumulative urine excretion showed that the FGF19^+^ mice displayed a similar response to water loading compared to control mice. This is in contrast with the increased urine output observed in FGF19+ mice when given free access to water (Figure 3A, 206 μL CI_95%_= [123; 290] for controls *vs* 433 μL CI_95%_= [330; 535] for FGF19, p=0.0013, and Figure 2C). These data suggest that FGF19 stimulates voluntary water intake, via the implication of the central nervous system.

**Figure 3:**
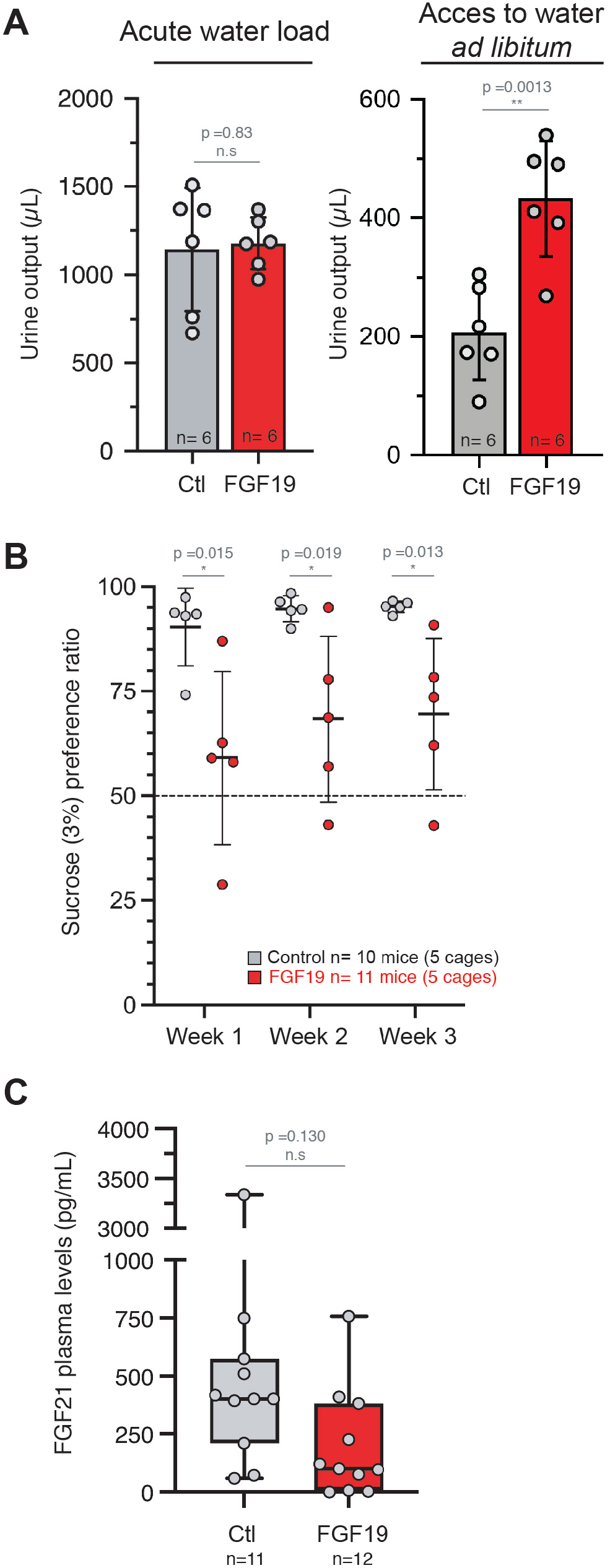
FGF19 effects on water homeostasis suggest central regulation. **A**. Urine output and urine osmolality 6h after acute water load or free acces to water. n= 6 mice per group. **B**. Sucrose (3%) preference ratio in a double-bottle experiment. Each dot represents a cage containing two to three animals. The total number of animals is indicated. **C**. FGF21 plasma levels determined by ELISA in control and FGF19 HGT mice. n= 11-12 mice per group. Data information: In (A-C): data are presented as mean ± SD. *P ≤0.05 (unpaired Student’s t-test). N.s: not significant.

The phenotype we report is reminiscent of what has been described for FGF21, where pharmacologic administration of the hormone also increased water consumption in mice (Camporez *et al*, 2013; Talukdar *et al*, 2016; Song *et al*, 2018; Turner *et al*, 2018). In a physiological context, FGF21 has been shown to stimulate water drinking in response to dehydration caused by metabolic stresses such as ketogenic diet or alcohol consumption (Song *et al*, 2018). Because FGF21 has also been involved in sugar appetence (Talukdar *et al*, 2016), we submitted control and FGF19 overexpressing mice to a two-bottle preference study between water and 3% sucrose. Strikingly, we observed the same phenotype as with FGF21, i.e. FGF19 overexpressing mice showed reduced appetence for 3% sucrose water (Figure 3B, at week 1: 90 % CI_95%_= [79; 102] for controls *vs* 59% CI_95%_= [33; 85] for FGF19, p=0.015). A possible mechanism for FGF19 to phenocopy the FGF21 effect is that the former increases the latter’s secretion. However, measurements of FGF21 plasmatic levels indicated that FGF19 expression had no effect on the FGF21 levels, ruling out an indirect effect (Figure 3C). Overall, we conclude that FGF19 effect on water homeostasis is central and independent of FGF21, however the exact mechanisms involved in this regulation remain to be determined. FGF21 impact on water homeostasis is mediated in part by activation of the neurons of the paraventricular nucleus in the hypothalamus (Song *et al*, 2018). FGF19 has been shown to exert effects on the central nervous system through FGFR1c/β-klotho signaling (Marcelin *et al*, 2014; Liu *et al*, 2018; Perry *et al*, 2015), therefore one possibility is that FGF19 activates FGFR1c in the hypothalamus. Another, and non-mutually exclusive, mechanistic explanation of FGF19 action is via modulation of the bile acid pool composition, since bile acids function as signaling molecules. Indeed, FGF19 is a potent inhibitor of bile acids synthesis by the liver and mice overexpressing FGF19 have modified bile acid pool composition, with increased FXR antagonists and decreased FXR agonists which in turn represses FXR activity (Gadaleta *et al*, 2018). It has been shown that FXR inactivation leads to a decrease in urine osmolality through the regulation of aquaporin water channel (aquaporin 2) in the renal collecting ducts (Zhang *et al*, 2014). Of note, one remarkable observation is the long-term phenotype after hepatic FGF19 expression: in contrast to effects of FGF21 on water intake after a single injection, we report a long-lasting diabetes insipidus phenotype.

### HCC patients with high circulating FGF19 have reduced natremia

To explore if the effects of FGF19 on water homeostasis have a clinical relevance, we considered pathological conditions associated with supra-physiological concentrations of FGF19. A subset of patients with hepatocellular carcinoma (HCC) have increased FGF19 serum levels (Maeda *et al*, 2019), and around 10% of tumors are characterized by the amplification of FGF19 genomic locus (Sawey *et al*, 2011; Schulze *et al*, 2015). In healthy subjects, FGF19 plasma levels range from 50 to 590 pg/mL (Lundåsen *et al*, 2006). To explore a potential effect on water homeostasis in patients with increased FGF19 circulating levels, we assayed FGF19 concentration in plasma in 173 HCC patients from the “Liver-pool Cohort”, held in the Montpellier University Hospital. Most patients were male (89%), had underlying cirrhosis (81%) and multifocal or advanced HCC stage (BCLC B, C or D stage, 73%). We defined three groups based on the result of the FGF19 ELISA assay (Figure 4A): “high FGF19” for the upper quartile ([FGF19] > 603 pg/mL, 43 patients), “intermediate FGF19” for the middle quartiles ([FGF19] between 218 pg/mL and 603 pg/mL, 87 patients) and “low FGF19” for the lower quartile ([FGF19] < 218 pg/mL, 43 patients). “High FGF19” group corresponded to supra-physiological concentrations based on the literature. Interestingly, natremia of the “high FGF19” group was significantly lower than the two other groups (Figure 4B, 140.1 mmol/L CI_95%_= [139.4; 140.8] for “low FGF19” *vs* 139.2 mmol/L CI_95%_= [138.5; 139.9] for “intermediate FGF19” *vs* 137.1 mmol/L CI_95%_= [136.2; 138] for “high FGF19”, p<0.0001). This result is suggestive of an imbalance in the regulation of the sodium and water homeostasis, potentially via an increased water consumption in a context of effective hypovolemia in this cohort of predominantly cirrhotic patients. Of interest, this effect remains significant when subjects under diuretic treatment or clinical ascites are excluded from the analysis (data not shown). Moreover, the natremia difference did not seem related to an impaired renal function in “FGF19 high” group (Figure 4C). Thus, our data from HCC patients with increased circulating FGF19 concentrations is consistent with an effect on water sodium balance in patients. Further work is required to establish if a reduction in natremia might have a prognostic value for FGF19-driven hepatocellular carcinoma. Finally, since FGF19 analogs are currently tested in different clinical trials, we suggest that it would be of great interest to analyze the newly discovered FGF19 activity, namely natremia and water intake, in response to these compounds.

**Figure 4:**
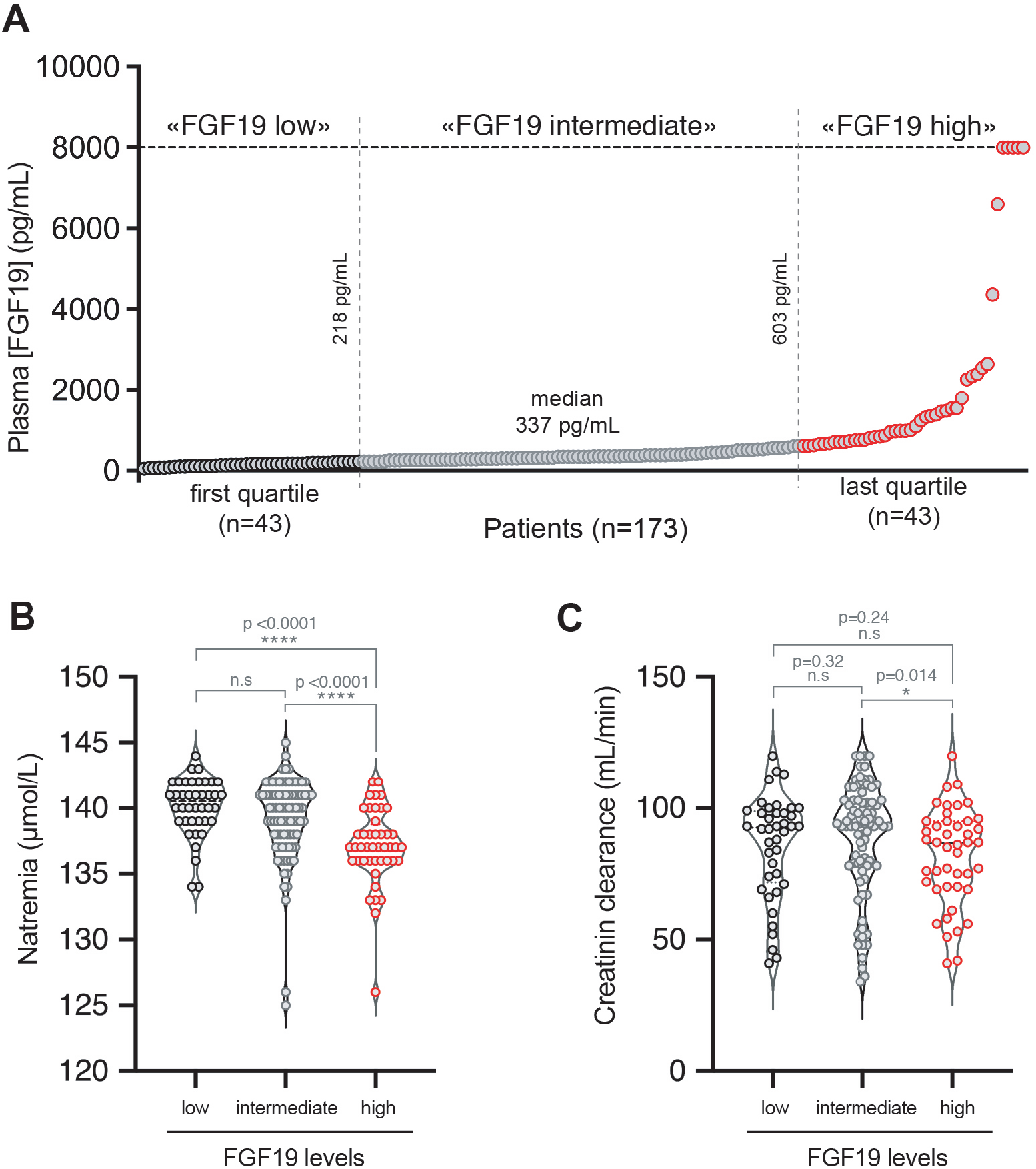
Increased FGF19 in hepatocellular carcinoma patients is associated with disturbances of the hydrosodic balance. **A**. Serum FGF19 distribution in a cohort of 173 patients with advanced HCC (FGF19 measured by ELISA). **B-C**. Violin-plots of natremia (B) and creatinin clearance (C) in patients of the cohort according to the plasma FGF19 concentration. Data information: * p ≤0.05, ** p<0.01, *** p<0.001, **** p<0.0001. N.s: not significant. Mann Whitney test significance is indicated.

## Material and Methods

### Vectors: cloning of FGF19/15

Plasmids constructs pSBbi-RN-FGF19 and pSBbi-BB-FGF15 were generated by cloning the human FGF19, amplified from Huh7 cDNA, and the murine FGF15, amplified from C57Bl/6J mouse ileum cDNA, into psBbi-RN and psBbi-BB plasmids (gifts from Eric Kowarz, Addgene plasmid #60519 and #60521, respectively), previously digested by the restriction enzyme SfiI. Primer sets for amplification of the inserts (with SfiI recognition site):

FGF19-SfiI-for-5’-ATCGGGCCTCTGAGGCCCAGGGAGGTGCCATGCGGA-3’

FGF19-SfiI-rev-5’-CGATGGCCTGACAGGCCGCCCCTGGCAGCAGTGAAGA-3’

FGF15-SfiI-for-5’-ATCGGGCCTCTGAGGCCCCCGAGGTGTCATGGCGAG-3’

FGF15-SfiI-rev-5’-CGATGGCCTGACAGGCCCGGAATCCTGTCATTTCTG-3’

### Animal experiments

All reported animal procedures were carried out in accordance with the rules of the French Institutional Animal Care and Use Committee and European Community Council (2010/63/EU). Animal studies were approved by institutional ethical committee (Comité d’éthique en expérimentation animale Languedoc-Roussillon (#36)) and by the Ministère de l’Enseignement Supérieur, de la Recherche et de l’Innovation (Apafis#10278-2018082809241782v1).

#### Hydrodynamic Gene Delivery

Hydrodynamic injections were performed in 6 to 8 week-old C57Bl/6J female (unless stated otherwise) mice, as described previously (Zhang *et al*, 1997; Liu *et al*, 1999). Briefly, 0.1 mL/g body weight of a solution of sterile saline containing plasmids of interest were injected into lateral tail vein over 8-10 s. psBBi-EF1a-FGF19-dTomato (12.5 μg), psBBi-EF1a-dTomato (12.5 μg), psBBi-EF1a-FGF15-TagBFP (12.5 μg) or psBBi-EF1a-TagBFP (12.5 μg) were injected together with sleeping beauty transposase SB100X (2.5 μg, ratio of 5:1). pCMV(CAT)T7-SB100 was a gift from Zsuzsanna Izsvak (Addgene plasmid #34879).

#### Allografts

C57Bl/6J female mice (Charles River) mice were anesthetized with intra-peritoneal injection of Xylazine-Ketamine mixture. After incision of abdominal wall and peritoneum, the left lateral lobe of the liver was pulled out of the mouse body. 5000 cells, resuspended in 5 μL of 25% Matrigel (BD) - PBS, were injected using a 10μL Hamilton syringe in the left lobe of the liver. After injection, liver was put back in normal position and the abdomen was sutured. Mice were monitored daily during 3 weeks, then euthanized before the tissues were collected and fixed following classical procedures.

Water consumption was measured by cages (n=2-5 mice) three times a week. For the double bottle assay, mice were acclimated to cages with two bottles containing water for one week. They were then given access to bottles with water containing 3% sucrose (Sigma-Aldrich #16104) or pure water. Water and water-3% sucrose intake was measured every two days. The position of bottles was changed every two days. Acute water loading was performed by an intraperitoneal injection of 2 ml of water. Mice were immediately placed in individual cages without access to food and water. Urine was collected hourly over 6 h.

### Blood samples for ELISA experiments

Mice were fasted for 4-6 hours then killed by anesthetic overdose with isoflurane. Plasma was collected after centrifugation. Plasma FGF19 and FGF21 were measured by ELISA, according to the manufacturers’ instructions: human FGF19 ELISA kit (Biovendor, RD191107200R), mouse FGF21 ELISA kit (Sigma-Aldrich, EZRMFGF21-26K).

### RNA isolation, qPCR

RNA was extracted from liver tissue and purified using RNeasy mini kit (Qiagen) according to manufacturer’s protocol. Reverse transcription of total RNA (1 μg) was done with QuantiTect Reverse Transcription kit (Qiagen), and cDNA quantified using LC Fast start DNA Master SYBR Green I Mix (Roche) with primers detailed below on LightCycler480 apparatus (Roche). Gene expression levels were normalized with hypoxanthine phosphoribosyltransferase (HPRT). Primer pairs used for qPCR: Hprt 5’-GCAGTACAGCCCCAAAATGG-3’ and 5’-GGTCCTTTTCACCAGCAAGCT-3’, FGF19 5’-CCAGATGGCTACAATGTGTACC-3’ and 5’-CAGCATGGGCAGGAAATGA-3’, Cyp7a1 5’-CTGCAACCTTCTGGAGCTTA-3’ and 5’-ATCTAGTACTGGCAGGTTGTTT-3’, Cyp8b1 5’-CCTGTTTCTGGGTCCTCTTATTC-3’ and 5’-TCTCCTCCATCACGCTGTC-3’.

### Human studies

A total of 173 patients from the prospective HCC “Liverpool” cohort conducted in the Montpellier University Hospital were included and analyzed retrospectively. All patients provided written consent for research at the time of their blood collection, in line with international regulations and ICH GCP (International Conference on Harmonization-Good Clinical Practice, Biobank Registration Number DC 2014-2328 AC 2014-2335). Montpellier University Hospital Institutional Review Board committee approved this study ((N° 2019_IRB-MTP_01-11). All the patients had FGF19 plasma concentration measured at the inclusion using human FGF19 ELISA kit (Biovendor, RD191107200R), according to the manufacturers’ instructions.

### Statistical Analyses

Data sets were tested with 2-tailed unpaired Student *t* tests or Mann-Whitney U tests, correlations were analyzed with Pearson’s χ^2^ test using Prism Software version 8 (GraphPad). Significant *P* values are shown as: \**P* <0.05, \*\**P* <0.01, \*\*\**P* <0.001, and \*\*\*\**P* <0.0001.

Comparisons of the mRNA expression levels between groups were assessed using Mann-Whitney U test. Spearman’s rank-order correlation was used to test the association between continuous variables. Univariate survival analysis was performed using Kaplan-Meier curve with log-rank test.

## Acknowledgments

We acknowledge Montpellier Biocampus facilities: the imaging facility (MRI), and the “Réseau d’Histologie Expérimentale de Montpellier” - RHEM facility, supported by SIRIC Montpellier Cancer (Grant INCa_Inserm_DGOS_12553), the European regional development foundation and the Occitanie region (FEDER-FSE 2014-2020 Languedoc Roussillon), for the histological analyses. We are grateful to zootechnicians of IGMM animal housing facility, Cédric Orféo, Eve Lasserre, Zoé Lebrere, for their work. We thank Emmanuel Vignal for the osmometer and Anne-Dominique Lajoix for allowing access to metabolic cages. We thank members of our lab for helpful discussions and comments. This work was funded by EVA-Plan cancer INSERM THE, and supported by SIRIC Montpellier Cancer Grant INCa_Inserm_DGOS_12553. The funders had no role in study design, data collection and analysis or publication process.

## Author contributions

JUB: Conceptualization, Methodology, Investigation, Writing-original draft, Writing-review & editing; CC: Conceptualization, Investigation, Writing-review & editing; LM: Investigation; GD: Investigation; IG: Investigation; AD: Resources, Investigation; TT: Resources; EA: Supervision, Funding acquisition, Writing-review & editing; UH: Supervision, Funding acquisition, Writing-review & editing; DG: Conceptualization, Investigation, Visualization, Supervision, Funding acquisition, Writing-original draft, Writing-review & editing.

